# Dissecting molecular regulatory mechanisms underlying noncoding susceptibility SNPs associated with 19 autoimmune diseases using multi-omics integrative analysis

**DOI:** 10.1101/871384

**Authors:** Xiao-Feng Chen, Min-Rui Guo, Yuan-Yuan Duan, Feng Jiang, Hao Wu, Shan-Shan Dong, Hlaing Nwe Thynn, Cong-Cong Liu, Lin Zhang, Yan Guo, Tie-Lin Yang

## Abstract

The genome-wide association studies (GWAS) have identified hundreds of susceptibility loci associated with autoimmune diseases. However, over 90% of risk variants are located in the noncoding regions, leading to great challenges in deciphering the underlying causal functional variants/genes and biological mechanisms. Previous studies focused on developing new scoring method to prioritize functional/disease-relevant variants. However, they principally incorporated annotation data across all cells/tissues while omitted the cell-specific or context-specific regulation. Moreover, limited analyses were performed to dissect the detailed molecular regulatory circuits linking functional GWAS variants to disease etiology. Here we devised a new analysis frame that incorporate hundreds of immune cell-specific multi-omics data to prioritize functional noncoding susceptibility SNPs with gene targets and further dissect their downstream molecular mechanisms and clinical applications for 19 autoimmune diseases. Most prioritized SNPs have genetic associations with transcription factors (TFs) binding, histone modification or chromatin accessibility, indicating their allelic regulatory roles on target genes. Their target genes were significantly enriched in immunologically related pathways and other immunologically related functions. We also detected long-range regulation on 90.7% of target genes including 132 ones exclusively regulated by distal SNPs (eg, *CD28, IL2RA*), which involves several potential key TFs (eg, CTCF), suggesting the important roles of long-range chromatin interaction in autoimmune diseases. Moreover, we identified hundreds of known or predicted druggable genes, and predicted some new potential drug targets for several autoimmune diseases, including two genes (*NFKB1, SH2B3*) with known drug indications on other diseases, highlighting their potential drug repurposing opportunities. In summary, our analyses may provide unique resource for future functional follow-up and drug application on autoimmune diseases, which are freely available at http://fngwas.online/.

**Author Summary:** Autoimmune diseases are groups of complex immune system disorders with high prevalence rates and high heritabilities. Previous studies have unraveled thousands of SNPs associated with different autoimmune diseases. However, it remains largely unknown on the molecular mechanisms underlying these genetic associations. Striking, over 90% of risk SNPs are located in the noncoding region. By leveraging multiple immune cell-specific multi-omics data across genomic, epigenetic, transcriptomic and 3D chromatin interaction information, we systematically analyzed the functional variants/genes and biological mechanisms underlying genetic association on 19 autoimmune diseases. We found that most functional SNPs may affect target gene expression through altering transcription factors (TFs) binding, histone modification or chromatin accessibility. Most target genes had known immunological functions. We detected prevailing long-range chromatin interaction linking distal functional SNPs to target genes. We also identified many known drug targets and predicted some new drug target genes for several autoimmune diseases, suggesting their potential clinical applications. All analysis results and tools are available online, which may provide unique resource for future functional follow-up and drug application. Our study may help reduce the gap between traditional genetic findings and biological mechanistically exploration of disease etiologies as well as clinical drug development.

## Introduction

Autoimmune diseases are groups of complex immune system disorders with high prevalence rates worldwide (4.5%) [1]. High heritabilities were observed on various autoimmune diseases (∼60%-90%) [2]. To date, genome-wide association studies (GWASs) have unraveled hundreds of susceptible loci associated with autoimmune diseases [3, 4], suggesting many functional genes involved in some key immunological pathways (eg. *MHC* gene clusters in antigen presentation, *TYK2* in cytokine signals) [5]. However, the true functional variants and target genes for the most of GWASs variants remain largely unknown [5], which might be mainly limited by two challenges. Firstly, the detected variants may be in linkage disequilibrium (LD) with causal functional SNPs without genotyping. Secondly, over 90% of GWASs variants are located in the uncultivated noncoding regions, complicating their functional interpretation.

In the past few years, many studies have integrated functional epigenetic data to predict function of noncoding SNPs. Many of these methods such as CADD [6], DeepSEA [7], GWAVA [8], FATHMM-MKL [9], ReMM [10] and FIRE [11], adopted machine learning algorithms to develop classifiers through integrating various annotations and labelled training data to distinguish potential functional/non-functional SNPs. However, the prior labelled training data may be inaccurate and impractical due to the current knowledge limitation in functional roles underlying noncoding SNPs. Some other methods like RegulomeDB [12], 3DSNP [13], GWAS4D [14], IW-Scoring [15], Eigen [16], and FunSeq2 [17] either directly combined various epigenetic/regulatory features to rank SNP functionality or adopted a weighted scoring scheme by considering the relative importance of each feature to assign SNP functionality scores. However, these approaches principally incorporated epigenetic or transcriptional annotation across all cells or tissues, while omitting the cell-specific or context-specific regulation. Besides, they mainly aimed to prioritize potential functional variants rather than dissect the downstream regulatory circuits linking functional variants to disease etiology. Autoimmune diseases associated variants are significantly enriched in blood cell-specific enhancers [18], implying that the integration of cell-specific functional data are required for dissecting molecular regulatory mechanisms underlying noncoding variants associated with autoimmune diseases.

The incorporation of cell-specific multi-omics data has remarkably accelerated the decryption of functional mechanisms underlying noncoding GWAS variants [19]. For example, we recently identified a functional SNP associated with two autoimmune diseases exerted allele-specific enhancer regulation on *IRF5* expression through long-rang loop formation [20]. Nevertheless, these studies primarily focused on one GWAS susceptibility loci on one disease, and only limited functional causal variants predisposing to autoimmune diseases have been validated [20]. The autoimmune diseases share substantial common susceptibility variants and immunopathology [21]. It is necessary and important to decipher the functions of GWAS noncoding variants systematically, which is helpful to accelerate the translation from GWASs findings into useful biological and clinical insights into autoimmune diseases.

To address these issues, we devised a new analysis frame to prioritize potential functional noncoding SNPs on 19 autoimmune diseases and further predicted their local and distal regulatory target genes using epigenetic, transcriptional and 3D chromatin interaction data across hundreds of blood immune cell types. Our analysis contains a new functional scoring method to prioritize functional autoimmune SNPs. We evaluated the performance of our functional scoring method by comparing it with other representative methods. We next explored potential molecular mechanisms underlying prioritized SNPs and analyzed the immunologically related function as well as potential clinical drug applications for predicted target genes. We also analyzed the roles of long-range chromatin interactions on autoimmune SNPs as well as potential key regulatory transcription factors (TFs). Finally, we developed an open web resource (http://fnGWAS.online/) and local analytical pipeline (https://github.com/xjtugenetics/fnGWAS).

## Results

### Prioritizing potential noncoding functional autoimmune SNPs

We collected 18,857 autoimmune noncoding tag SNPs predisposing to 19 distinct autoimmune diseases (*P* < 5×10^−8^) from multiple resources (Table S1). LD analysis retained 51,594 noncoding tags and LD expanded (*r*^2^ > 0.8) SNPs in 333 genome-wide significant loci (autoimmune positive SNPs). We next collected 26,922,878 background SNPs in all 333 loci, and collected 47,131,427 negative SNPs beyond these loci. To explore potential key epigenetic regulatory features for autoimmune diseases, we compared 606 epigenetic data annotation across 47 blood immune cell types between all autoimmune positive SNPs and background SNPs. Previous studies had suggested that the autoimmune causal SNPs are significantly enriched in blood-cell specific enhancer marks [18]. Consistently, we found that autoimmune positive SNPs are significantly higher enriched for 347 active epigenetic features (FC > 1, *P* < 0.05/606) across 40 blood immune cell types within four epigenetic categories, including 9 DHSs, 75 active histone modifications (H3K4me1, H3K4me2, H3K4me3, H3k27ac and H3K9ac), 167 active genomic segmentations (HMM-15, marked as active transcription or enhancer) and 96 TFBS (Figure 1a and Table S2). To evaluate the functionality of all positive SNPs, we devised a new epigenetic functional scoring approach (flowchart shown in Figure S1) using fold enrichment of all 347 significant epigenetic features across four epigenetic categories as weight. By comparing functional score of each positive SNPs with scoring distribution of negative SNPs, we prioritized 15,314 SNPs associated with 19 autoimmune diseases with functionality support on at least one epigenetic category (Figure 1b-c and Table S3).

**Figure 1.**
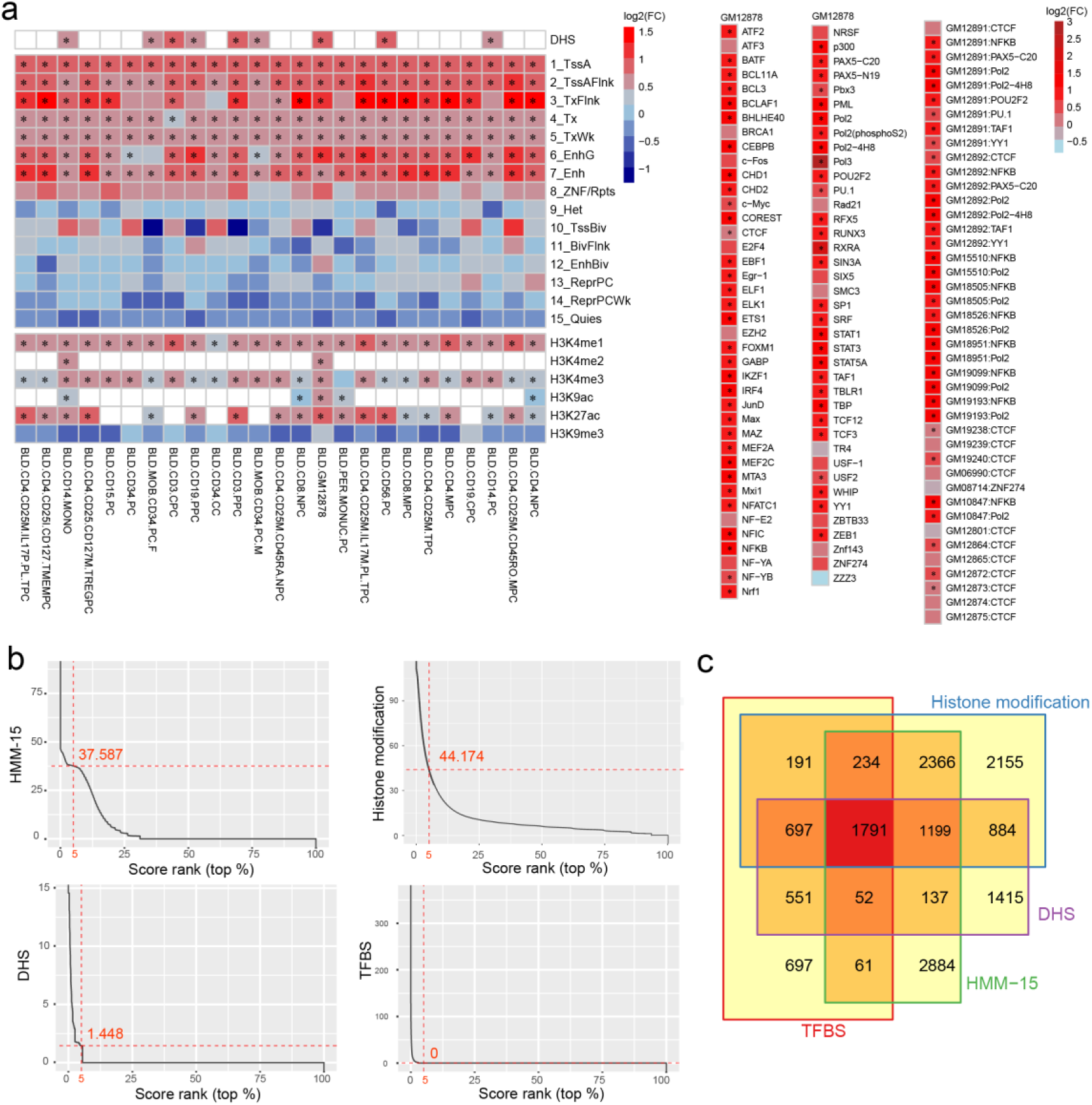
Epigenetic functional scoring for autoimmune SNPs. (a) Heatmap showing epigenetic feature enrichment analysis on 606 epigenetic data from 47 blood cell types across four epigenetic groups (Left: DHS, HMM-15, histone modification. Right: TFBS) between all autoimmune positive SNPs and background SNPs. FC: fold enrichment on each feature comparing autoimmune positive SNPs with background SNPs. Red color represents feature with higher enrichment in autoimmune positive SNPs (Log_2_FC > 0). All significant and active features (Bonferroni adjusted *P* < 0.05, FC > 1) selected for SNP scoring were marked with asterisk. Enrichment analysis was performed using Fisher’s exact test. (c) Ranking plot for scores of all autoimmune negative SNPs within four epigenetic categories, with red dashed line represents top 5% ranked value. (d) Venn diagram showing count of autoimmune SNPs with functionality support in each of four epigenetic categories by comparing the scoring value with top 5% ranked value of all negative autoimmune SNPs in (a). See also Figure S1.

### Integrative prediction of potential causal target genes on prioritized SNPs

To explore potential regulatory targets for 15,314 prioritized SNPs, we integrated both cis-QTL association, 3D chromatin interaction and colocalization analysis from over 30 blood cell types (Table S4). We predicted 367 high-confident target genes regulated by 4,272 prioritized functional SNPs (Table S5 and S6), which had both cis-QTL, chromatin interaction and colocalization evidence (PP4 > 0.8).

### Functional SNPs are significantly enriched in allele-specific motif and local molecular QTLs

The functional SNPs might perturb allelic enhancer activities through mediating several intermediate molecular-level traits, such as bQTL [22], hQTL [23], dsQTL [24] or caQTL [25] (Figure 2a). To explore potential allelic regulatory mechanisms linking 4,272 prioritized autoimmune SNPs to predicted gene targets, we firstly performed motif analysis, and detected allele-specific binding motif on 2,603 SNPs (Figure 2a). We further analyzed multiple molecular QTL association (Table S7) for autoimmune SNPs, and identified 592 SNPs associated with several intermediate-level molecular traits (Table S8), including 143 bQTL SNPs preferentially binding to special allele on 5 TFs (JunD, NF-kB, PU.1, Pou2f1, Stat1) in LCLs, 303 caQTL or dsQTL SNPs affecting chromatin accessibility in either naive or stimulus-specific macrophages (n = 157) or CD4+ T cells (n = 24) or LCLs (n = 182), as well as 230 hQTL SNPs affecting chromatin modification on either H3K4me1 (n = 63), H3K4me3 (n = 83) or H3K27ac (n = 127) in LCLs (Figure 2a). Further analysis revealed significant enrichment for all molecular QTL association (bQTL, dsQTL, caQTL and hQTL) on prioritized functional SNPs compared with all autoimmune SNPs (FC = 1.9 ∼ 7.4, *P* < 0.05, Figure 2b-e), implying their extensive regulatory roles. We also detected weak while significant enrichment for allele-specific binding motif on prioritized functional SNPs in comparison with all autoimmune SNPs (FC = 1.05, *P* = 1.59×10^−14^, Figure not shown), further supporting their important regulatory roles. Together, these analyses suggested potential allelic regulatory mechanisms underlying 66.0% of prioritized autoimmune functional SNPs.

**Figure 2.**
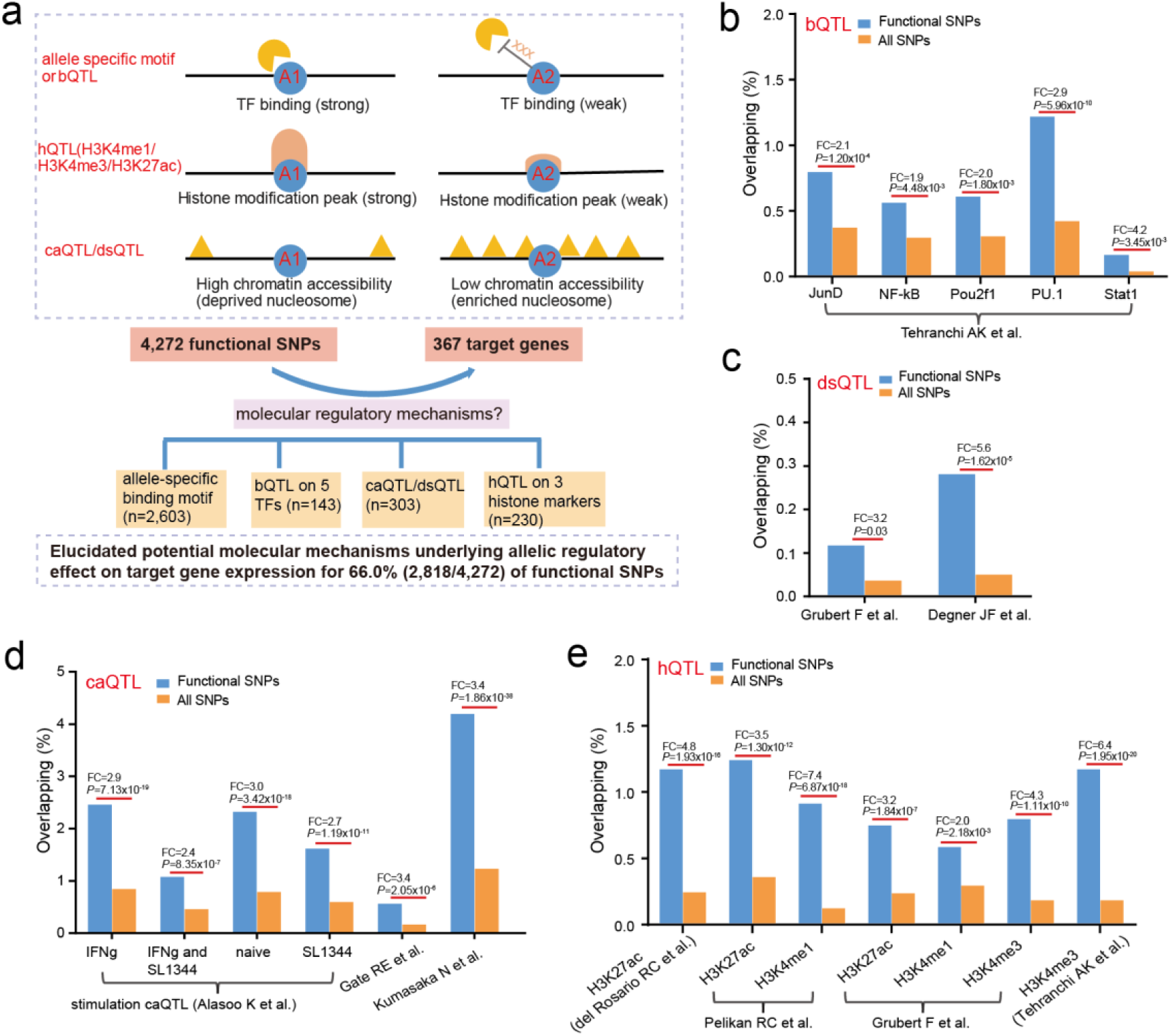
Dissecting allelic regulatory mechanisms underlying functional SNPs. (a) Schematic showing several potential molecular-level regulatory mechanisms underlying functional autoimmune SNPs (upper) and summary of corresponding SNP counts (bottom). (b-e) Functional enrichment for each collected molecular QTL data on functional SNPs compared with all positive autoimmune SNPs. Fisher’s exact test was performed in b-e, with fold enrichment and *P*-value shown.

### Epigenetic functional scoring improves prioritizing functional autoimmune SNPs compared with other methods

To further assess the performance of our epigenetic functional scoring, we compared the functional support on multiple immune-cell associated regulatory evidence between SNPs prioritized by our method and other five functional scoring methods [11-15]. Table S15 summarized the main characteristics between our method and other scoring methods (see discussion for comparison in detail). To ensure fair comparison of methods performance, we extracted top-ranked SNPs under different functionality support by our method with corresponding equal or approximately equal counts of top-ranked SNPs from other methods, which resulted in comparison with two methods under all functionality support and another three methods under selected functionality supports (Figure 3a).

**Figure 3.**
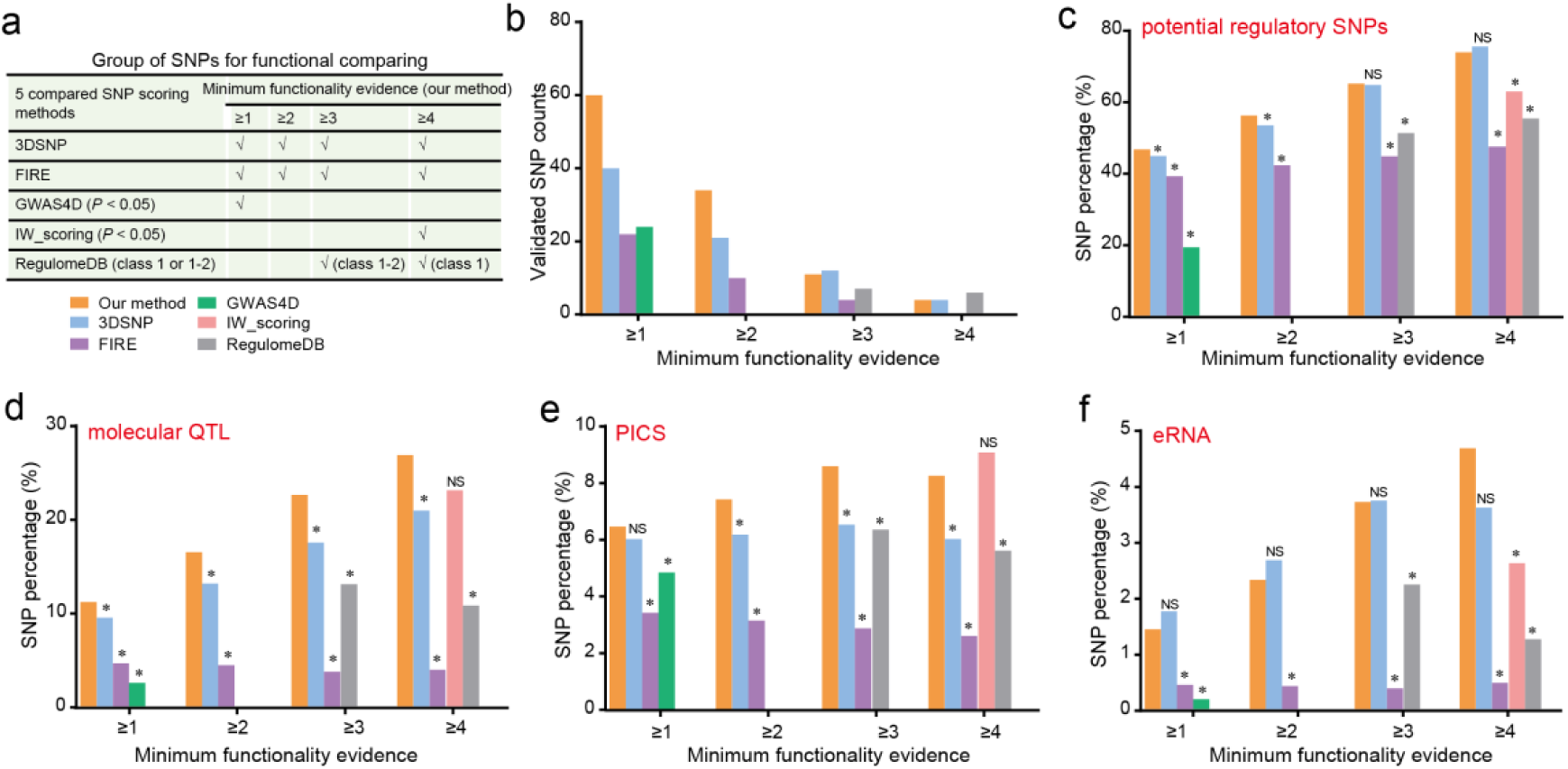
Comparing epigenetic functional scoring with other methods. (a) Description of top-scored SNPs sets from each method for functional comparing in b-f, which are marked by different colors (bottom). (b) Comparison of experimentally validated functional SNPs between our method and other five methods from a high-throughput screen assay in mononuclear cells [26]. (c-f) Comparison of percentage of annotated SNPs within different regulatory evidence between our method and other five methods, including (c) potential regulatory SNPs with predicted target gene by combining cis-QTL and chromatin interaction analysis, (d) potential functional SNPs with significant molecular QTL (bQTL, hQTL, dsQTL or caQTL) association, (e) casual autoimmune associated SNPs identified by PICS approach [18], and (f) potential regulatory SNPs within eRNA detected from IBD patients [28]. Fisher’s exact test was performed in c-f with asterisk represented significant higher enrichment on our method (FC > 1, *P* < 0.05). NS, not significant. See also Figure S2.

We firstly compared experimentally validated regulatory SNPs in mononuclear cells [26], and detected substantially more validated SNPs by our method compared with either FIRE, GWAS4D or IW-Scoring (Figure 3b). We also detected much more validated SNPs compared with 3DSNP under the first two functionality support and comparable validated SNPs compared with RegulomeDB (Figure 3b). Consistent results were found on experimentally validated regulatory SNPs in two no-immune cell types (K562, HepG2) [27], in which our method had substantially more validated SNPs compared with all four methods except for 3DSNP (Figure S2a-b). In comparison with 3DSNP, we identified comparable experimentally validated SNPs in two no-immune cell types (Figure S2a-b) while substantially more validated ones in the mononuclear cell (Figure 3b), implying the potential outperformance of our method in prioritizing immune cell specific regulatory SNPs. We next compared potential regulatory SNPs under multiple immune-related functional evidence (potential regulatory SNPs associated with gene expression, SNPs with molecular association, causal SNPs identified by PICS approach [18], eRNA SNPs from IBD patients [28]). We found that our prioritized SNPs was significantly higher enriched for nearly all functional evidence compared with all five methods (Fisher’s exact test, FC > 1, *P* < 0.05, Figure 3c-f). We also detected much higher percentage of molecular QTL SNPs or eRNA SNPs for our prioritized SNPs under the highest functionality support compared with either IW-Scoring (FC = 1.2, *P* = 0.08, Figure 3d) or 3DSNP (FC = 1.3, *P* = 0.06, Figure 3f) although not significant. Collectively, these analyses support the outperformance in prioritizing functional autoimmune SNPs by our epigenetic functional scoring method over other mentioned comparable methods.

### Target genes are significantly enriched in immunologically related functions

To evaluate the immunologically related functions on predicted target genes on 4,272 prioritized SNPs, we collected multiple immune-relevant functional datasets. We identified 181/367 highly-supported potential immunological genes (Figure 4a and Table S9), including 171 genes participated in immunologically related pathways, 25 genes whose knockdown in mouse could display abnormal immune system phenotypes from IMPC portal [29], as well as 23 genes associated with Mendelian disorders with immunology-related clinical symptoms from the OMIM database. We further analyzed other suggestive immune-relevant functions for predicted target genes, and detected functional support for nearly all (365/367) target genes (Figure 4a and Table S9), including 358 genes expressed on 20 blood immune cell types (RPKM > 1), 39 genes with tissue-specifically expression on blood as determined by TSEA approach (pSI < 0.01) [30], 191 immune system diseases associated genes collected from the DisGeNET database [31], as well as 201 genes showed causal relationship with autoimmune diseases as implemented the SMR analysis (FDR < 0.05, *P*_HEIDI_ > 0.05, Table S10) [32]. Collectively, these data suggested potential immunological function for most gene targets, which may suggest new mechanistic insights into autoimmune disease etiologies.

**Figure 4.**
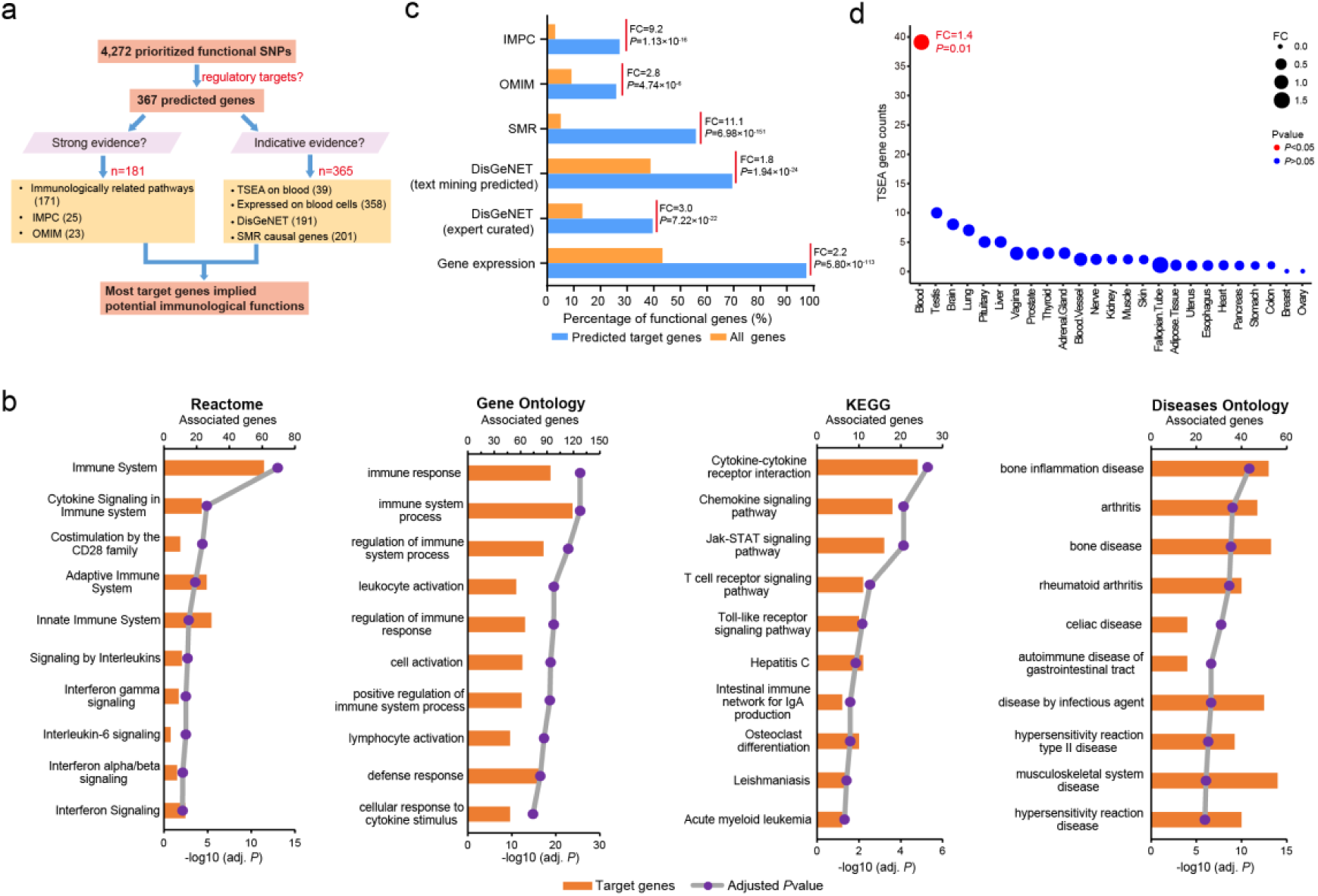
Immunological function analysis for predicted target genes. (a) Summary of multiple immunologically related functions for predicted target genes. (b) Top 10 significant biological pathways on predicted target genes. Both *P*-value (line chart) and gene counts (bar chart) are shown. (c) Functional enrichment for potential immunologically related gene set with different functional evidence between predicted target genes and whole genome genes. Enrichment analysis was performed using Fisher’s exact test. (d) Tissue Specific Expression Analysis (TSEA) for predicted target genes on 25 diverse tissues, with dot size representing gene counts and dot color indicating significance level (*P*-value) using Fisher’s exact test. Only one significant (*P* < 0.05) tissue (blood) was detected.

To further verify the immunological roles for predicted target genes, we performed functional enrichment analysis. We found that the predicted target genes are significantly enriched in multiple immunologically related pathways (Figure 4b). We also detected significant enrichment for other immunologically related genes from different functional datasets (IMPC, OMIM, DisGeNET) and SMR causal genes and expressed genes on blood cell types on predicted target genes (Fisher’s exact test, FC: 1.8 ∼ 11.1, *P*: 4.74×10^−6^ ∼ 6.96×10^−151^, Figure 4c). We further compared tissue-specific expression from TSEA [30] on 25 distinct cell types, and detected exclusively significant higher enrichment for blood tissue (Fisher’s exact test, FC = 1.4, *P* = 0.01, Figure 4d) on predicted target genes, which also showed the largest number of tissue-specifically expressed genes (Figure 4d). Altogether, these analyses revealed extensive enrichment of immunologically related functions for target genes, supporting the credibility of our target gene prediction.

### Prevailing long-range regulation linking functional autoimmune SNPs to distal target genes

Among 367 prioritized target genes, we detected larger amount of distal genes (n = 333, regulated by distal functional SNPs) compared with local genes (n = 235, regulated by functional SNPs located within target gene promoter), including 132 distal genes exclusively regulated by distal functional SNPs (Figure 5a). These exclusive distal genes included many known immunologic genes, such as *CD37, CD28, IL7, IL12RB1* or *IL2RA*, indicating the important roles of long-range regulation on autoimmune diseases. We further analyzed all 7,221 SNP-gene regulatory pairs, and detected predominantly distal pairs (87.87%) compared with local ones (Figure 5b). Interesting, the distal SNPs residing within local genes are more likely to regulate the distal target genes compared with their directly located genes (64.8% vs 17.89%, Figure 5b). We also analyzed the distance between all distal regulatory pairs, and found that the vast amount of distal SNP-gene regulatory pairs (66.5%) are located more than 50 kb away (mean distance: 105.4 kb, Figure 5c), further underscoring the important roles of chromatin looping on autoimmune diseases.

**Figure 5.**
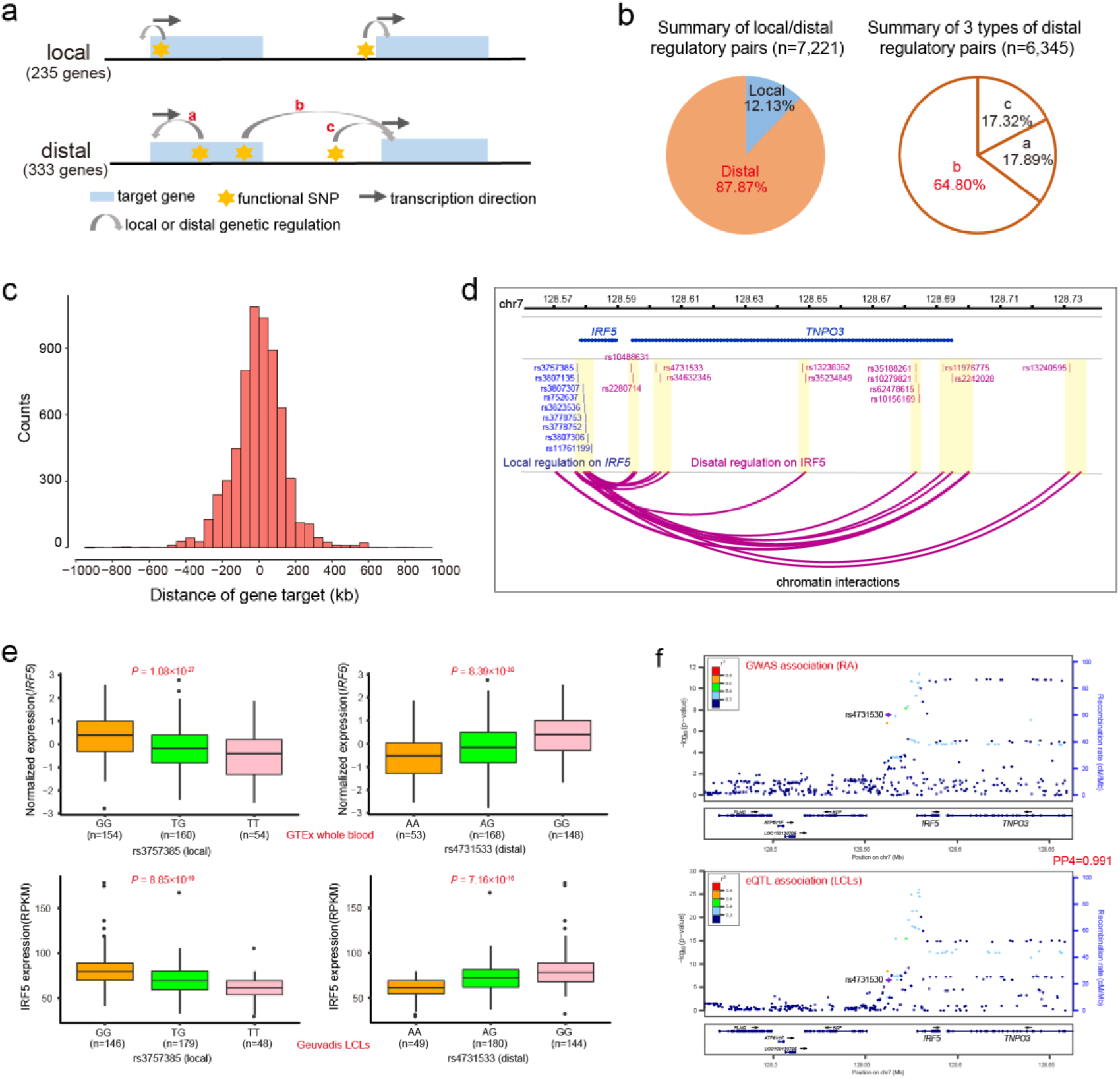
Prevailing long-range regulation linking functional SNPs to distal gene targets. (a) Schematic showing different regulatory models underlying prioritized functional autoimmune SNPs and gene targets. (b) Pie chart showing comparison between local and distal regulatory pairs (left), as well as between 3 types of distal regulatory pairs (right) in a. (c) Counts of SNP-gene pairs at different distance (kb). (d-f) *IRF5*-*TNPO3* region example showing that multiple functional autoimmune SNPs located within other gene regulated distal target gene expression via long-range chromatin interactions. The regulatory evidence including both (d) chromatin interactions, (e) cis-QTL association (one local example SNP and another distal example SNP were shown) and (f) colocalization between GWAS association on RA and *IRF5* cis-eQTL association in LCLs. Genomic annotation and chromatin interaction at *IRF5* locus were visualized using WashU Epigenome Browser.

The prevailing long-range regulation may indicate that, for many functional noncoding autoimmune SNPs, their located or directly mapped genes might not be the direct regulatory target genes. Among 3,139 prioritized functional SNPs within gene region (intergenic SNPs excluded), we found that 67.1% of SNPs exclusively regulated 239 distal effect genes instead of directly located local genes. We show one such example in Figure 5d-f, in which multidimensional evidence (cis-QTLs, 3D chromatin interactions and colocalization) supported that multiple functional SNPs within *TNPO3* or in the intergenic region near *TNPO3* could regulate distal *IRF5* expression through long-range chromatin interactions. The *IRF5* was also locally regulated by several functional SNPs within *IRF5* promoter region (Figure 5d-f). The immunological roles of *TNPO3* is largely unknown. In contrast, *IRF5* is a well-known immunological gene with crucial roles in autoimmune etiology [33], thus providing plausible mechanistic insights linking GWAS risk SNPs at *IRF5*-*TNPO3* locus to autoimmune pathogenies [20].

### Distal autoimmune genetic regulatory network may be mediated by several key TFs

To identify potential functional TFs mediating genetic regulation for autoimmune diseases, we compared allele-specific motif occupying between 4,272 functional SNPs and all autoimmune SNPs. We identified 29/366 nominally significant (Fisher’s exact test, *P* < 0.05) motif TFs with higher enrichment for functional SNPs (Figure 5a and Table S11). To explore potential regulatory targets on prioritized TFs, we considered three possible TF-gene regulatory models (Figure 6b), including (1) local model: TFs directly bind to target gene promoter to mediate gene expression, distal model: TFs bind to distal enhancers to regulate target gene expression via long-range chromatin interactions, and (3) indirect model: the TFs regulate target gene expression through mediating other regulatory TFs in trans manner. We found that most of our predicted target genes (72.8%, 267/367) could be regulated by these 29 TFs (Figure 6b-c and Table S11), with predominant distal model (n = 218) compared with either local model (n = 102) or indirect model (n = 112). Moreover, CTCF had the most regulatory target genes (Table S11), consistent with its known role in facilitating long-range chromatin looping [34]. Further analysis revealed that all 29 TFs had more distal regulatory target genes compared with local genes (Figure 6d), and 25 of them involved potential immunological functions (Figure 6e), implying their broad roles in distal genetic regulation on autoimmune diseases. We further analyzed the sharing of gene targets between different TFs, and detected 22 TFs sharing all 267/267 target genes with all 7 other rest TFs (Figure 6f), indicating their potential central regulatory roles. Together, these analyses suggested several possible key regulatory TFs mediating distal genetic regulatory networks on autoimmune diseases.

**Figure 6.**
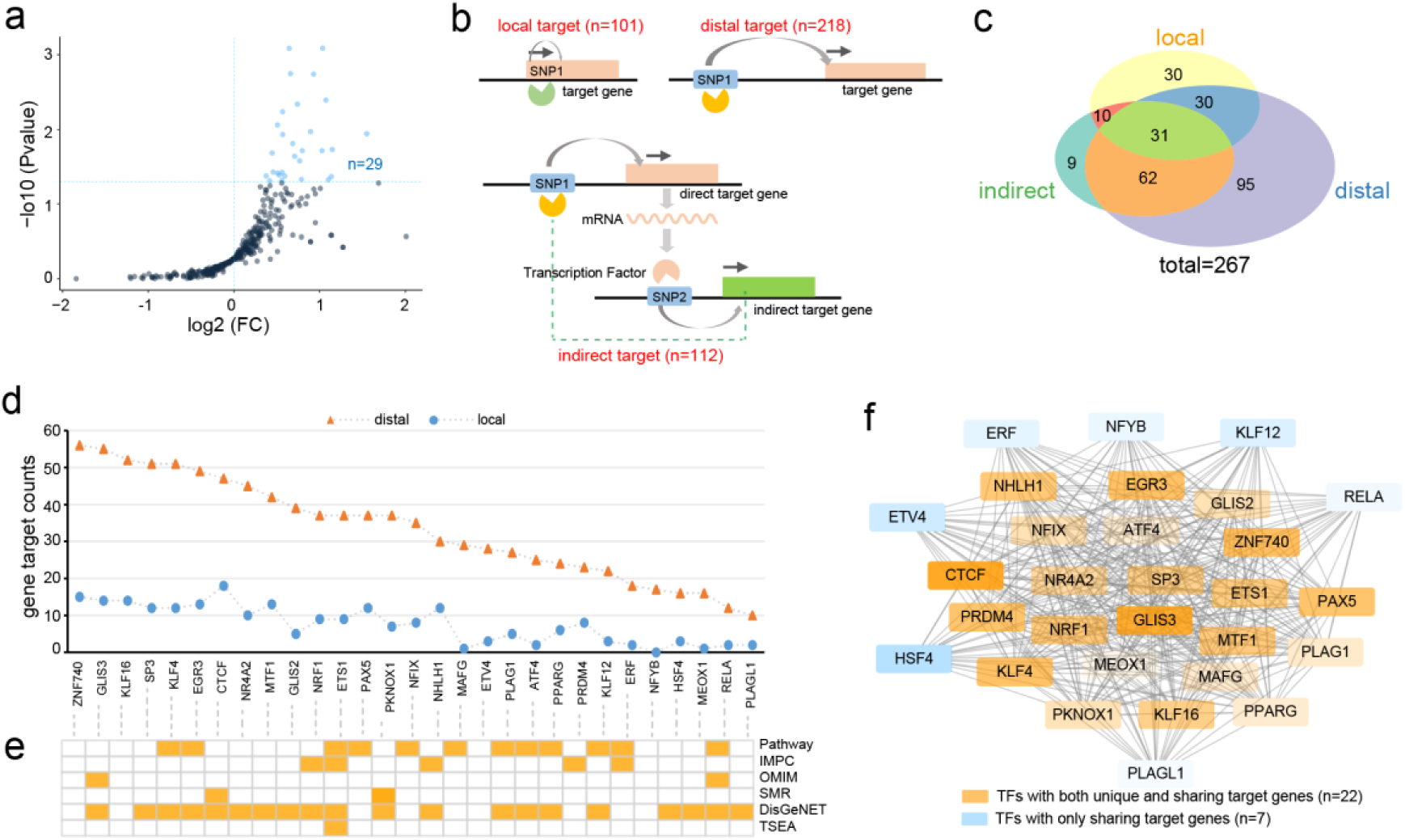
Identifying key TFs mediating autoimmune genetic regulatory network. (a) Scatter plot showing fold enrichment (FC) and significance level for enrichment in 366 predicted motif TFs between prioritized functional SNPs and all autoimmune SNPs. Nominally significantly higher enriched TFs (FC > 1, *P* < 0.05) are marked in blue. (b) Schematic showing three TF-target gene regulatory models. The gray arrow indicates SNP-target gene interaction. (c) Venn diagram showing counts of three types of target genes on significant TFs in (b). (d) Comparison between distal and local target genes on significant TFs. (e) Annotated immunological functions on significant TFs. (f) Pervasive sharing of regulatory target genes between different significant TFs. The orange rectangle represented 22 TFs shared all target genes with the rest (blue) of TFs, which might indicate their central regulatory roles. The transparency indicated counts of regulatory target genes, with CTCF mediated the most target genes (n = 113).

### Analyzing potential clinical applications on target genes

To explore potential clinical implications on predicted target genes, we firstly investigated all approved or experimental drug targets with known indications. We identified 80 genes targeted by drugs with known clinical indications on either autoimmune diseases (n = 41) or other immunologically related diseases (eg, allergies, infections or inflammations, n = 45) or other diseases (n = 57) (Figure 7a-b, Table S12), implying the extensive therapeutic implications on predicted target genes. The identified drug target genes showed pervasive shared drug indications, with 62.2 % of genes targeted for other immunologically related diseases and 42.1 % of genes targeted for other diseases also shared targeted indications for autoimmune diseases (Figure 7b and Table S12), indicating potential pleiotropic therapeutic-effect among drug targets. Except for known drug target genes, we also identified 190 potential druggable genes, including 118 ones without known drug target indications (Figure 7a and Table S13). In comparison with all genome genes, our predicted target genes are significantly more enriched in both known drug target genes (Fisher’s exact test, FC = 4.3, *P* = 2.93×10^−28^) and predicted druggable genes (Fisher’s exact test, FC = 3.5, *P* = 6.94 ×10^−61^) (Figure 7c), further supporting the potential important clinical implications on them.

**Figure 7.**
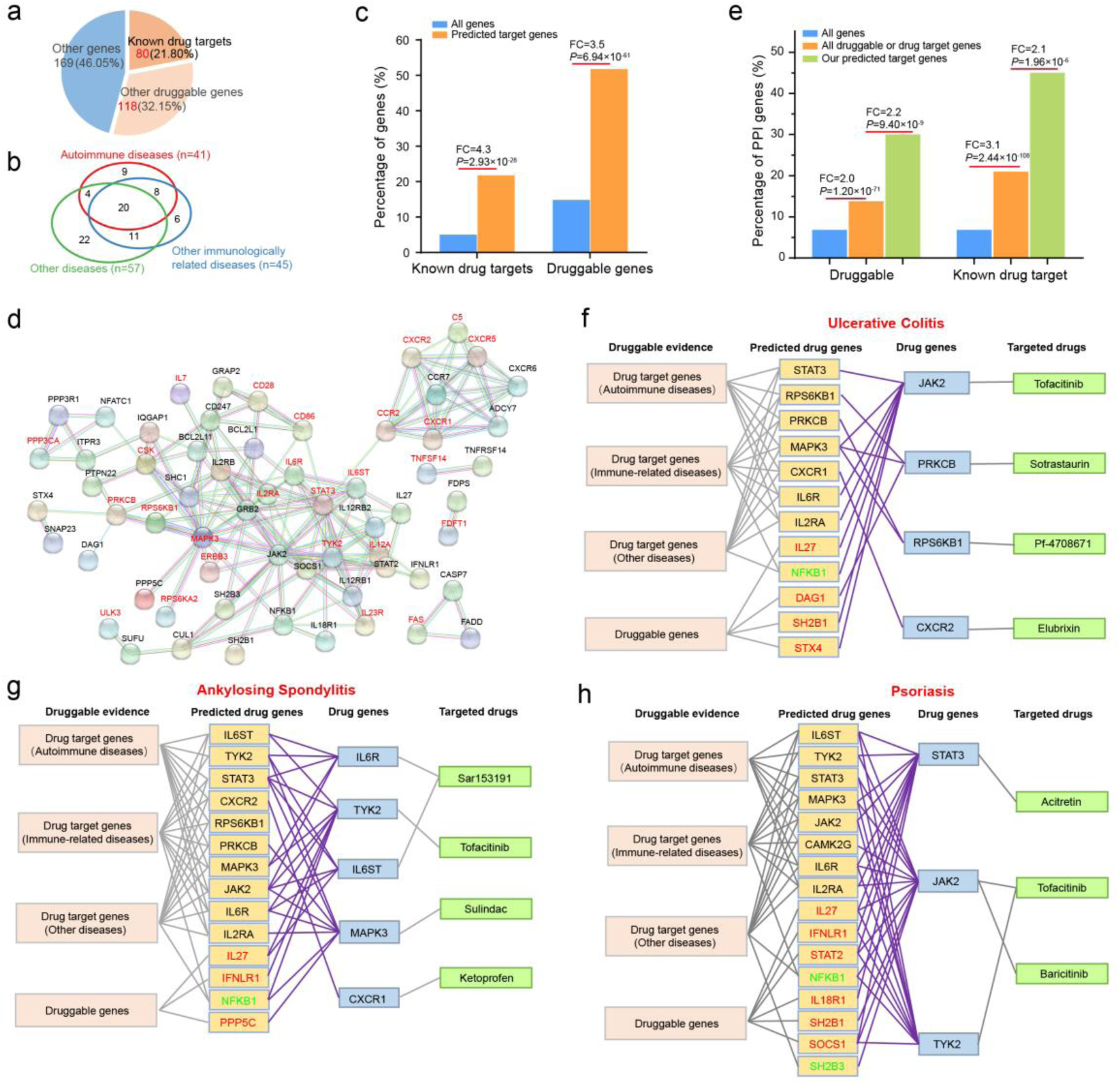
Predicted new drug targets with potential repurposing opportunities for three autoimmune diseases. (a) Pie chart showing gene count and percentage among predicted target genes for either known drug targets with indications or predicted druggable genes or others. (b) Venn diagram showing sharing counts of drug target genes with indications on either autoimmune diseases, other immunologically related diseases or other diseases (Table S12 for detail). (c) Functional enrichment analysis for either known drug target or predicted druggable genes on our predicted target genes compared with all genome genes using Fisher’s exact test. (d) PPI (score > 0.9) between autoimmune-drug target genes (marked in red) and other drug target or druggable genes. PPI plot was from STRING database by querying online. (e) Functional enrichment analysis showing percentage of genes showing strong PPI (score > 0.9) with autoimmune-drug target genes on either predicted druggable genes or known drug target genes. The comparison was performed between our predicted target genes (marked in green) and all druggable or drug target genes (marked in orange), as well as between all druggable genes or drug targets and all genome genes (marked in blue) using Fisher’s exact test, respectively. (f-h) Predicted new candidate drug targets on three autoimmune diseases. The orange rectangle shows predicted new drug genes, with genes with known indications on other autoimmune or non-autoimmune diseases marked in black or green and genes without known drug target indications marked in red. See also Figure S3 and S4.

Consistent with the observed pleiotropic indications among drug target genes (Figure 7b), we found extensive disease association sharing for both autoimmune drug target genes and other drug target or druggable genes (Figure S3a-b), which may suggest new potential opportunities for drug repurposing on autoimmune diseases from other non-autoimmune drug target or druggable genes. To explore the functional relevance between known autoimmune-drug genes and other genes, we firstly analyzed their shared biological pathways. We found that the vast majority (36/41) of autoimmune-drug genes shared the same immunologically related pathways with 68.9 % (131/190) of drug target or druggable genes (Figure S3c), implying their intimately functional connectivity. We further performed protein-protein interaction (PPI) analysis, and detected strong PPI (interaction score > 0.9) between 63.4 % (26/41) of autoimmune-drug genes and 31 other known drug target or druggable genes (Figure 7d), indicating the pervasive regulatory relevance between known autoimmune-drug target and other genes. This was further supported by enrichment analysis, in which 41 autoimmune-drug genes showed significantly higher PPI with either predicted druggable genes (FC = 2.0, *P* = 1.20×10^−71^) or known drug target genes (FC = 3.1, *P* = 2.44×10^−108^) compared with whole genome genes (Fisher’s exact test, Figure 7e). Besides, when restricted PPI targets of autoimmune-drug genes to our predicted gene targets, we found significantly higher PPI on our predicted target genes compared with either all predicted druggable genes (FC = 2.2, *P* = 9.40×10^−9^) or all known drug target genes (FC = 2.1, *P* = 1.96×10^−6^) (Fisher’s exact test, Figure 7e). Based on these analyses above, it is reasonable to assume that incorporating both GWAS genetic regulation and protein interaction network could help prioritize new potential drug target genes for autoimmune diseases. We prioritized 25 new candidate drug target genes for seven autoimmune diseases (Figure 7f-h and S5, Table S14), which showed both strong PPI with known drug target genes and genetic regulation associated with the same autoimmune disease. Among 25 prioritized genes, we found 14 genes with known indications on other autoimmune diseases as well as 2 genes (*NFKB1, SH2B3*) with indications on other diseases (Figure 7f-h and S4). The rest 9 genes had no indications while with druggable evidence, including 4 genes (*DAG1, IL27, STX4*, SH2B1) predicted targeted for ulcerative colitis and 3 genes (*IL27, IFNLR1, PPP5C*) predicted targeted for ankylosing spondylitis as well as 6 genes (*IFNLR1, SOCS1, IL27, STAT2, IL18R1, SH2B1*) predicted targeted for psoriasis (Figure 7f-h). Together, our analysis not only prioritized some new promising drug targets for future drug exploration, but also suggested some known drug targets (*NFKB1, SH2B3*) that could be exploited for future drug repurposing on autoimmune diseases.

### Open web application and local pipeline

To facilitate quick search for interested SNP(s) or gene(s) prioritized by our integrative analysis, we developed an open website (http://fngwas.online/) collecting comprehensive resources including functional scores on all noncoding autoimmune SNPs, regulatory target genes on prioritized functional SNPs, immunologically related functions for predicted target genes, clinical drug applications for target genes as well as regulatory mechanisms underlying functional SNPs. We also provided precomputed functional analysis results across whole genome SNPs/genes for bulk downloading (http://fngwas.online/download.php), which included functional scores and predicted allelic regulatory mechanisms underlying all autosomal noncoding SNPs as well as multiple disease-relevant function and drug target analysis for all genome genes. To further expand the potential application of our analytical frame on other complex diseases/traits, we also developed packaged local pipeline named fnGWAS (dissecting the functionality of noncoding GWAS SNPs, workflow shown in Figure S5), which could be run on any local Linux server with user-definable annotation data and parameters (https://github.com/xjtugenetics/fnGWAS).

## Discussion

The majority of autoimmune susceptibility SNPs are located in the noncoding region. It remains challenging to pinpoint the causal SNPs and functional genes to decipher the underlying biological mechanisms. In this study, we systematically evaluated the molecular mechanisms underlying noncoding susceptibility SNPs associated with 19 autoimmune diseases, through combining functional SNPs prioritizing, target gene prediction, allelic regulatory mechanisms analysis, gene function annotation as well as drug application exploration. We found predominant long-range chromatin interaction linking functional SNPs to distal target genes, which may be mediated by several key TFs including CTCF. Particularly, we detected extensive regulatory roles underlying prioritized functional SNPs, as well as broad immunological functions and clinical drug applications on predicted target genes. We also developed open website and analytical pipeline. We hope that our systematic analyses may be helpful for future experimental follow-up as well as clinical exploitation of drug repurposing on autoimmune diseases.

We have previously integrated epigenetic features for known disease-associated SNPs to predict novel susceptibility loci for complex diseases [35-38]. In this study, we developed a new improved epigenetic functional scoring method to prioritize functional autoimmune SNPs through incorporating hundreds of immune cell-specific active epigenetic information. Some other comparable scoring methods are also developed, such as 3DSNP [13], FIRE [11], GWAS4D [14], IW-Scoring [15] or RegulomeDB [12]. Compared with these approaches, one distinct feature of our method was the integrating of immune cell-specific epigenetic information (Table S15), which might provide better evaluation for disease-specific functional autoimmune SNPs. Another feature of our analysis frame is the comprehensive functional evaluation on multiple regulatory levels spanning SNP functional scoring, gene target prediction, gene function analysis and gene clinical application analysis, as well as SNP regulatory mechanisms analysis (Table S15). Indeed, the integration of cell-specific epigenetic annotation has been proved highly successful for prioritizing functional GWAS SNPs validated by experimental assays in many recent studies [39, 40]. Our analysis revealed that the top-ranked autoimmune SNPs prioritized by our method are significantly higher enriched in multiple blood immune cell associated regulatory elements compared with other methods, implying the outperformance of our method. We anticipated that future incorporation of more other cell-specific or context-specific epigenetic information could help identify functional SNPs associated with other complex diseases/traits.

Recent studies have shown that considerable noncoding GWAS SNPs could regulate target genes through long-range loop formation [41-43], providing unprecedented new mechanical insights underlying GWAS disease association. Consistently, our analysis revealed prevailing long-range regulation linking functional autoimmune SNPs to distal target genes, suggesting the important roles of chromatin interactions for autoimmune diseases. Our analysis also suggested that many functional SNPs within local gene could regulate distal target gene expression, including vast amounts of functional SNPs within local gene promoter. One underlying mechanism hypothesis was that gene promoter could also act as enhancer which was termed Epromoter to regulate distal gene expression [44], which was consistent with our recent findings that one functional autoimmune risk SNP within *TNPO3* promoter could independently regulate distal *IRF5* expression via long-range loop formation [20]. We also identified several potential key regulatory TFs with significant enrichment in functional autoimmune SNPs, including CTCF. CTCF is well-known for its regulatory roles for mediating enhancer-promoter interaction in chromatin loop formation [34], and played essential roles in late B-cell differentiation [45]. In line with the prevailing long-range genetic regulation detected for autoimmune diseases, we found predominant distal regulatory genes compared with local ones for all enriched TFs, indicating their potential roles in mediating distal genetic regulatory network for autoimmune diseases. Future functional assays are needed to decipher their precise regulatory mechanisms.

The past fruitful GWAS findings have remarkably accelerated the translation of new drug clinical utilities [4]. The drug targets with human genetic evidence of disease association are twice as likely to lead to approved drugs [46]. Consistently, we found that our predicted autoimmune target genes are significantly more enriched in both known drug target genes and druggable genes compared with whole genome genes, supporting the potential important clinical implications on disease effecter genes. A previous GWAS study has incorporated PPI with 98 annotated RA risk genes to predict new drug targets, and highlighted *CDK6* and *CDK4* as promising candidates [47]. The incorporation of functional genomic and immune-related annotations as well as PPI has been demonstrated successfully in prioritizing potential drug target on immune-related traits [48]. Consistently, our study integrated both genetic association and PPI, and prioritized 25 candidate drug target genes on 7 autoimmune diseases, including many genes (16/25) with known indications on autoimmune diseases or other diseases. The drug repurposing strategies have shed light on many new promising therapeutic opportunities for autoimmune diseases, such as the dopaminergic drug for multiple sclerosis [49] or Fibrate for treating for primary biliary cirrhosis [50]. Our results may provide important clues for future clinical drug repurposing on autoimmune diseases. For example, we predicted *IL2RA* to be a potential new drug target for ankylosing spondylitis. *IL2RA* is targeted by several known drugs (eg. HuMax-TAC) with indications on autoimmune diabetes and has known roles in the pathogenesis of autoimmunity [5]. Besides, we found that *IL2RA* was regulated by several functional SNPs associated with ankylosing spondylitis. Collectively, these evidence suggest the potential drug repurposing opportunity of *IL2RA* on ankylosing spondylitis.

In conclusion, we performed comprehensive functional genetic analysis for 19 autoimmune diseases. We hope that our unique resource may help accelerate the translation from GWASs findings into biologically and clinically useful insights underlying autoimmune diseases pathogenies.

## Materials and Methods

### Autoimmune SNPs collection

We collected SNPs associated with 19 autoimmune diseases [alopecia areata (AA), ankylosing spondylitis (AS), autoimmune thyroid disease (ATD), celiac disease (CEL), Crohn’s disease (CRO), IgE and allergic sensitization (IGE), inflammatory bowel disease (IBD), juvenile idiopathic arthritis (JIA), multiple sclerosis (MS), narcolepsy (NAR), primary biliary cirrhosis (PBC), primary sclerosing cholangitis (PSC), psoriasis (PSO), rheumatoid arthritis (RA), systemic lupus erythematosus (SLE), systemic scleroderma (SSc), type1 diabetes (T1D), ulcerative colitis (UC), and vitiligo (VIT)] from multiple resources, including the GWAS Catalog [3], the ImmunoBase (https://www.immunobase.org/) and other public studies [51, 52]. All databases were visited in March 2019 and summarized in Table S1. For SNPs achieved genome-wide significance reported in European ancestry (*P* < 5×10^−8^), any coding or splicing SNPs annotated by ANNOVAR [53] using GENCODE v19 reference data were removed. We further excluded SNPs within the major histocompatibility complex locus (MHC, https://www.ncbi.nlm.nih.gov/grc/human/regions/MHC?asm=GRCh37.p13) due to the complex LD patterns. The filtered SNPs were selected as autoimmune tag SNPs.

### LD analysis, positive, background and negative SNPs definition

LD analysis for autoimmune tag SNPs was conducted using PLINK v1.90 [54] in European samples from 1000 genome v3 genotype data [55], with maximum distance for *r*^2^ calculation set as 1M. Genome-wide significant loci were defined as merged unique regions surrounding 1M of any filtered noncoding tag SNPs with overlapping MHC region truncated. We extracted noncoding tags and LD expanded (*r*^2^ > 0.8) SNPs within genome-wide significant loci as positive SNPs and all noncoding SNPs in these loci as background SNPs. We collected 41,377 susceptible SNPs with ID record in the 1000 genome v3 genotype data [55] from GWAS Catalog (visited in March 2019). All other noncoding SNPs beyond genome-wide significant loci and beyond MHC region with low LD (*r*^2^ < 0.1) with the GWAS catalog susceptible SNPs were selected as negative SNPs.

### Epigenetic functional scoring

#### Epigenetic features selection

We collected 606 epigenetic data (called peak region) on 47 blood cell types from Roadmap [56] and ENCODE Project [57]. Four different epigenetic categories of data were incorporated for SNP annotation, including 15 chromatin states (HMM-15), histone modification, DNase I hypersensitive sites (DHS) and transcription factor binding sites (TFBS). One epigenetic feature represents one epigenetic annotation in one cell type (eg, H3K4me1 in GM12878). SNPs were labeled as annotated or unannotated on each epigenetic feature by analyzing their overlapping with selected feature using bedtools v2.25.0 [58]. We performed enrichment analysis for each epigenetic feature by comparing counts of annotated positive SNPs and background SNPs using chi-square test. All epigenetic features with significantly higher enrichment for positive SNPs compared with background SNPs (Fold enrichment > 1, Bonferroni adjusted *P* < 0.05) were selected for the following epigenetic scoring. The fold enrichment (FC) is defined as:

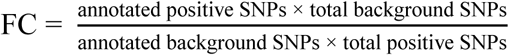

#### Functional scoring

Based on our previous epigenetic enrichment approach [37, 38], we developed a new cell-specific epigenetic weighted scoring method to evaluate the functionality for all noncoding autoimmune positive SNPs (flowchart shown in Figure S1). For each epigenetic category (HMM-15, histone modification, DHS, TFBS), we adopted an accumulative quantitative score system using fold enrichment of selected significant features within each category as weight, separately, which is defined as follows:

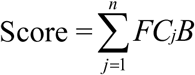

Where j denotes particular feature (1 ≤ j ≤ n) among each epigenetic category (assuming n total features), B indicates whether the tested positive SNP was annotated (B = 1) or unannotated (B = 0) on feature j. Therefore, we can get four independent functional scores across four different epigenetic categories for each tested SNP. For each epigenetic category, we further scored for all negative SNPs to build null distribution, and prioritized any positive SNPs with score higher than the top 5% ranked score value of all negative SNPs as potential functional. Finally, any positive SNPs with functionality support in at least one epigenetic category were determined as potential functional.

### Predicting target genes for prioritized functional SNPs

#### Cis-QTL analysis

We examined the cis-quantitative trait loci (cis-QTLs) association between prioritized noncoding SNPs and all nearby genes in 1M region. We collected 12 cis-eQTL and 2 cis-pQTL data over 20 blood immune cell types from 13 different published studies (Table S4). For pQTL data from the INTERVAL study [59], we extracted all cis-pQTL (1M surrounding gene TSS) pairs and transformed the protein ID to gene symbol ID using the UniProt online tools. For any full QTL dataset without multiple testing corrections, we adjusted original *P* using false discovery rate (FDR) method. All significant QTL results with probe/gene level FDR <5% validated by at leaste two different datasets were retained.

#### Three-dimensional (3D) chromatin interaction analysis

All SNP-gene pairs with cis-QTL associations were divided into either local (SNPs within target gene promoter (1KB surrounding TSS)) or distal (SNPs beyond target gene promoter). We collected chromatin interaction assay (5C, in situ Hi-C, capture Hi-C, HiChIP, ChIA-PET) and predicted chromatin interaction data (IM-PET, PreSTIGE, PHM) on multiple blood immune cell types from 11 different studies (Table S4). To validate the long-range regulation between distal SNP-gene pairs, the 3D chromatin interactions between prioritized SNP and gene transcript promoter region (GENCODE v19) were examined using bedtools v2.25.0 [58]. The integration of cis-QTLs and 3D chromatin interactions might better identify causal regulatory effect at GWAS loci by diminishing the potential accidental overlapping with QTLs for GWAS SNPs [60]. All distal SNP-gene pairs with chromatin interaction evidence from at leaste two different datasets were retained.

#### Co-localization analysis

To validate the potential causal genetic regulatory effect for filtered local or distal target genes, we employed two complementary methods [61, 62] to assess whether the detected GWAS signal and cis-QTL association shared the same causal variant. For 16 GWAS summary and 7 full QTL datasets available (Table S1 and S4), we employed the Coloc [61] method using coloc R package for Co-localization analysis. The Coloc method [61] adopted a Bayesian statistical test using summary-level data to estimate five posterior probabilities: no association with either GWAS or QTL (PP0), association with GWAS while not with QTL (PP1), association with QTL while not with GWAS (PP2), association with GWAS and QTL while with two independent SNPs (PP3), association with both GWAS and QTL with one shared causal SNP (PP4). We defined 100-KB region surrounding each GWAS index SNP (*P* < 5×10^−8^) and tested for co-localization with any overlapping QTL genes. For all curated GWAS and QTL datasets (including datasets with no full summary-level data available, Table S1 and S4), we also employed another adapted Coloc method named PICCOLO [62] for Co-localization analysis. The PICCOLO [62] estimates the colocalization of GWAS and QTL PICS (Probabilistic Identification of Causal SNPs) [18] credible set using reported lead SNPs and *P-*value. The PICS was a fine-mapping algorithm to estimate each SNP’s probability of being causal at a given locus [18]. We performed PICCOLO analysis as described using piccolo R package by Tachmazidou et al. [63]. Briefly, we firstly estimated the PICS credible set for each lead GWAS index SNP and each top QTL SNP using pics.download and then performed colocalization analysis using pics.coloc.lite with default parameter. For both Coloc and PICCOLO, any genes with both PP4 greater than 80% and significant QTL association with prioritized SNPs from at least two cis-QTL datasets were considered to support the co-localization.

#### Local and distal target gene prediction

We predicted local or distal target gene on prioritized SNPs using different strategies. For local ones, any genes with both cis-QTLs assocaition and colocalization evidence were prioritized to be potential target genes. For distal ones, any genes with multidimensional evidence including cis-QTLs assocaition, 3D chromatin interaction and colocalization were considered to be potential target genes.

### Deciphering allelic regulatory mechanisms underlying prioritized SNPs

#### Allele-specific motif analysis

We analyzed the allelic effect of prioritized functional SNPs on transcription factor binding motifs using FIMO from MEME Suite toolkit (v4.11.0) [64] with default parameters and TF motifs available from 5 public motif databases, including JASPAR (2018 version) [65], HOCOMOCO (v11) [66], SwissRegulon [67], Transfac and Jolma2013 [68]. To identify potential functional motifs, we focused motif search on TF genes with high expression in at least one of the 20 blood immune cells from Roadmap [56] or DICE [69] (RPKM >1). The allele-specific binding motifs predicted by at least two different datasets were retained.

#### Molecular QTL analysis

We collected different molecular QTL data in multiple blood cell types from 8 studies (Table S7), including transcription factor binding quantitative trait loci (bQTL) on five immune-relevant TFs (NF-κB, PU.1, Stat1, JunD, and Pou2f1), histone modification quantitative trait loci (hQTL) (H3K4me1/H3K4me3/H3K27ac), DNase-I hypersensitivity quantitative trait loci (dsQTL) and chromatin accessibility quantitative trait loci (caQTL). For all QTL datasets, the tested SNP and molecular peak (TF binding sites or ChIP-Seq peaks) pairs could be divided into either local (SNP located within molecular peak) or distal ones (SNP located beyond molecular peak). We retained significant association results between prioritized functional SNPs and local molecular peaks which passed multiple testing corrections (FDR < 0.1).

### Comparison with other functional scoring methods

#### Curation of top-ranked SNPs

We compared our epigenetic functional scoring with five other functional scoring methods, including 3DSNP [13], FIRE [11], GWAS4D [14], IW-Scoring [15] and RegulomeDB [12]. The IW-Scoring [15] integrated eleven commonly used scoring methods to assign SNP a combined significance level (*P*-value) and outperformed any single method. We therefore did not compare our method with these eleven methods. Functional scores of all autoimmune positive SNPs from these methods were collected from online database in March 2019. We extracted prioritized autoimmune SNPs by our method under four different minimum functionality evidence (⩾4, ⩾3, ⩾2, ⩾1, n = 1,791 ∼ 15,331), and extracted equivalent or approximately equivalent top-ranked SNPs by other five methods for functional comparison. (1) Since both 3DSNP and FIRE adopted the quantitative scoring system, we selected those top scoring ranked SNPs equal to our prioritized SNPs under different minimum evidence (⩾4, ⩾3, ⩾2, ⩾1) for functional comparison, respectively. (2) The GWAS4D calculated combined regulatory probability (*P*-value) for examined variants by jointly considering cell type-specific regulatory potential and cell type-free composite score. We retained significant SNPs on GM12878 (*P <* 0.05, n = 16,868) for comparison with our prioritized SNPs under at least one functional evidence (⩾1, n = 15,314), which had approximately equal SNP counts. (3) Similarly, we selected significant SNPs (*P <* 0.05, n = 341) by IW-scoring for functional comparison with our prioritized SNPs under at least four evidence (⩾4, n = 1,791), which had the closest SNP counts. (4) The RegulomeDB adopted a category based scoring system (class from 1-7, with lower rank means higher functional support). We extracted SNPs ranked within class 1 (n = 1,958) or within class 1-2 (n = 3,575) for functional comparison with our prioritized SNPs under at least three (⩾3, n = 3,973) or four evidence (⩾4, n = 1,791), respectively, which had the closest SNP counts.

#### Functional enrichment comparison

For collected functional SNPs set from each methods, we firstly compared their experimentally validated SNPs count in three cell types (blood mononuclear cells, K562 and HepG2) from two recent high-throughput screen reports [26, 27]. We next compared their functionality enrichment on multiple regulatory data support using Fisher’s exact test, including (1) SNPs with predicted local or distal target genes by integrating cis-QTL and chromatin interaction analysis on over 30 blood immune cell types (Table S4), (2) SNPs annotated with molecular QTL (bQTL, hQTL, dsQTL and caQTL) on multiple blood immune cell types (Table S7), (3) reported causal SNPs associated with 16 autoimmune diseases (AA, AS, ATD, CEL, CRO, JIA, MS, PBC, PSC, PSO, RA, SLE, SSC, T1D, UC, VIT) prioritized by the PICS approach [18], and (4) SNPs annotated with enhancer RNA (eRNA) from IBD patient samples [28].

### Exploring immunologically related functions for predicted target genes

#### Pathway analysis and functional genes curation

We performed biological pathway enrichment analysis (including Gene Ontology [GO], Kyoto Encyclopedia of Genes and Genomes [KEGG], Disease Ontology [DO] and Reactome pathway) for all predicted gene targets using clusterProfiler R package with default parameter [70], except that setting use_internal_data = TURE for KEGG enrichment analysis to enable online query from latest KEGG data. To identify potential immunologically related genes, we manually curated immunologically related biological pathways from all annotated terms on predicted target genes. We also collected immunologically related genes from other public datasets, including the International Mouse Phenotyping Consortium (IMPC) portal (http://www.mousephenotype.org/, release-9.2), the Online Mendelian Inheritance in Man (OMIM) database (https://www.omim.org/), and the DisGeNET database (http://www.disgenet.org/home/, v6.0, expert curated or text mining predicted genes) [31]. All dataset were downloaded or queried online in May 2019.

#### Gene expression and tissue-specific expression analysis

We collected gene expression data on 5 blood immune cell types (CD4 memory, CD4 naïve, Mobilized CD34, Peripheral blood mononuclear, GM12878) from Roadmap [56] and 15 primary immune cells types from the DICE project (http://dice-database.org/) [69]. Gene expression was measured by RPKM (reads per kilobase per million mapped reads). We collected the gene lists with tissue-specific expression (as based on a specificity index threshold [pSI], pSI < 0.01) in 25 broad GTEx tissue types from report by Wells et al. [30].

#### SMR analysis

We analyzed the causal relationship between predicted target genes and autoimmune diseases risk using 16 GWAS summary and 7 QTL summary data (Table S1 and S4) by the summary data–based Mendelian randomization (SMR) approach [32]. We ran SMR (v0.712) with default parameters. LD correlations between SNPs were estimated from 6,743 unrelated European samples from the Atherosclerosis Risk in Communities (ARIC) data (dbGap: phs000280.v3.p1.c1) [71] with one of each pair of individuals with a SNP-derived relatedness estimate of > 0.025 suggested by GCTA (v1.91) [72] randomly removed. Gene-disease pairs passed both multi-SNP-based SMR test (FDR adjusted *P*_SMR_ < 0.05) and heterogeneity test by HEIDI (*P*_HEIDI_ > 0.05) were considered to be potential causal.

### Regulatory TF analysis

We performed enrichment analysis for all allele-specific binding TFs on functional autoimmune SNPs by comparing annotated functional SNPs with all positive autoimmune SNPs using Fisher’s exact test. For each TF with significant higher enrichment on autoimmune SNPs (*P* < 0.05, FC > 1), we assigned the predicted regulatory targets of its binding SNPs as its direct regulatory target genes. The TF-gene regulatory network was visualized by Cytoscape V3.4 (http://www.cytoscape.org/).

### Drug target and drug repurposing analysis

#### Curation of drug target genes

Clinically approved or experimental drug target genes with known indications were obtained from 3 different databases, including the DrugBank database (https://www.drugbank.ca/, v5.1.2) [73], the Therapeutic target database (TTD, 2018 updated) [74] and Open Targets database [75]. All three drug databases were queried in March 2019. For TTD dataset, we translated the UniProt protein ID into corresponding gene symbol ID using UniProt online tools. All drug indications were manually classified into autoimmune diseases, immunologically related diseases (allergies, infections, inflammations, rejection, immune system diseases and hematologic malignancies) or other diseases.

#### Curation of druggable genes

We collected potentially druggable genes from either DGIdb (www.dgidb.org, v3.0.2) [76], Pharos (https://pharos.nih.gov/idg/targets) [77] or report by Finan et al. [78]. We queried DGIdb and Pharos in March 2019. The DGIdb organized druggable genome under two classes, including over 35 validated or predicted drug-gene interaction types from 20 disparate sources, and 39 gene categories associated with druggability. The Pharos classified all targets into 4 groups by characterizing the degree to which they are not studied (labeled Tdark) or studied (labeled Tbio, Tchem or Tclin). The studied targets from Pharos were retained. Any gene targets with druggability evidence from at least two resources were prioritized as potentially druggable.

#### Predicting new potential drug target genes

For all annotated drug target or druggable genes, we analyzed protein-protein interaction (PPI) between these genes and all other genes. PPI was queried online from the STRING database (https://string-db.org/) in June 2019 with only high-confident interacted pairs (interaction score > 0.9) retained. By leveraging both PPI and upstream autoimmune diseases regulatory information, we can prioritize new potential drug target gene A or for particular disease B by filtering: (1) A has strong PPI (interaction score > 0.9) with any drug target gene C which had known indication on autoimmune disease B, (2) Both A and C are regulated by upstream functional SNPs predisposing to autoimmune disease B, (3) A is either known drug target gene or predicted druggable gene. The predicted genes with known indication on other disease might suggest new potential drug repurposing opportunities.

### Functional enrichment analysis

Functional enrichment for all collected immune-relevant functional datasets (IMPC, OMIM, SMR, DisGeNET, TSEA, gene expression, drug target) on predicted target genes was analyzed by comparing annotated target genes with whole genome genes in each dataset using Fisher’s exact test. Functional enrichment for immune-cell associated regulatory data (motif, molecular QTL) on prioritized functional SNPs was analyzed by comparing annotated functional SNPs with all positive autoimmune SNPs using Fisher’s exact test.

## Data availability

All analysis results are free for searching online or bulk downloading at http://fngwas.online.

Analysis pipeline scripts are available at https://github.com/xjtugenetics/fnGWAS.

## Description of Supplemental Data

Supplemental Data contains 5 supplementary figures and 15 supplementary tables.

## Acknowledgments

We thank the ARIC Communities study. We obtained ARIC data through dbGaP authorized access at https://dbgap.ncbi.nlm.nih.gov/aa/wga.cgi?page=login with the accession number of data phs000280.v3.p1.c1.We are grateful to Dr. Ruihua Jing for the constructive discussions of manuscript preparation.

## Supplementary Figures Legends

**Figure S1. Workflow of epigenetic functional scoring**

The top panel shows definition for positive, background and negative autoimmune SNPs for the following epigenetic functional scoring. Any coding, spicing or MHC region SNPs were removed. The middle panel shows the process for functional scoring. FC: fold enrichment. Epigenetic data in 47 blood immune cell types across four epigenetic categories (HMM-15, histone modification, DHS, TFBS) are used for enrichment analysis using Fisher’s exact test. M1-M4 denotes annotated or unannotated positive/background SNPs count on each epigenetic feature. A1-A4 denotes four epigenetic categories with m1-m4 significant enriched features for scoring. The bottom panel shows how to determine functionality support for each positive SNP. Each SNP had four scores (n1-n4) across four epigenetic groups, which were further compared with 5% top ranked score value of all negative SNPs (S1-S4) to determine its functionality support. Relative to Figure 1.

**Figure S2. Comparing epigenetic functional scoring with other methods using experimentally validated regulatory SNPs**

Comparison of experimentally validated functional SNPs between our epigenetic functional scoring and other five methods from high-throughput screen assay in HepG2 (a) and K562 (b) cells [27]. Relative to Figure 3.

**Figure S3. Prevailing sharing of genetic disease-association and biological pathways on drug target genes**

(a-b) Count of (a) autoimmune drug target genes or (b) other drug target and predicted druggable genes associated with paired autoimmune diseases, with genes associated with individual disease shown in diagonal line. Disease association on gene targets are derived from their upstream functional SNPs. (c) Counts of shared immunological related pathways between 41 known autoimmune-drug target genes (row) and all 198 drug target or druggable genes (column). Pathways were manually curated from all annotated biological terms (GO, KEGG, DO, Reactome) on predicted target genes (Table S9). Relative to Figure 7.

**Figure S4. Predicted new potential drug targets for four autoimmune diseases**

The yellow rectangle shows predicted new drug genes for four autoimmune diseases which had strong PPI with known drug target genes (blue). All predicted drug genes had known indications on other autoimmune diseases or non-autoimmune diseases. Relative to Figure 7.

**Figure S5. Flowchart of fnGWAS pipeline**

The blue rectangle summarized five main analysis steps of fnGWAS pipeline, with aim for each step shown (Step 1-5). For each analysis step, the input data (represented by cylinder) and simplified example summarized output result (represented by yellow table) are shown, respectively. By default, fnGWAS begins with an epigenetic functional scoring pipeline (Step1) using all susceptible SNPs associated with any interested diseases/traits as input, which outputs functional scores and functionality support for all positive SNPs (see detailed workflow for step 1 in Figure S1). Target gene prediction were then employed for all positive SNPs with functionality support (Step 2). Downstream functional analysis were then performed predicted target genes and their regulatory functional SNPs (Step 3-5). Alternatively, each step of fnGWAS can be run independently, which support any user-defined input data. The whole pipeline including input annotation data are free available at https://github.com/xjtugenetics/fnGWAS or http://fngwas.online/download.php.

**Supplementary Tables**

**Table S1**. Summary of datasets for autoimmune SNPs collection

**Table S2**. Significantly enriched active epigenetic features selected for epigenetic functional scoring

**Table S3**. Epigenetic functional scores on all positive autoimmune SNPs

Table S4. Summary of cis-QTLs and chromatin interactions datasets for target gene prediction

**Table S5**. Predicted regulatory target genes on prioritized potential functional SNPs

**Table S6**. Colocalization between GWAS and cis-QTL association for predicted target genes

**Table S7**. Summary of intermediate molecular QTL datasets

**Table S8**. Potential molecular regulatory mechanisms underlying functional autoimmune SNPs

**Table S9**. Summary of immunologically related functions for target genes

**Table S10**. Potential causal autoimmune target genes identified by SMR

**Table S11**. Regulatory target genes and immunologically related functions for significantly enriched TFs

**Table S12**. known drug target genes with clinical indications

**Table S13**. Prioritized candidate druggable genes

**Table S14**. Predicted new potential drug target or drug repurposing genes on autoimmune diseases

**Table S15**. Comparison between fnGWAS and other representative scoring approaches

